# Influence of traffic and weather on carcass persistence time of small wildlife on roads

**DOI:** 10.1101/2023.06.01.543226

**Authors:** Annaëlle Bénard, Christophe Bonenfant, Thierry Lengagne

## Abstract

The rapidly expanding road network threatens the persistence of many terrestrial species through habitat loss, fragmentation, chemical, light and noise pollution and mortality associated with wildlife-vehicle collisions. Roadkill monitoring under-estimate actual collision numbers as the time during which roadkill carcasses remain visible on the road is often shorter than the frequency of road monitoring. By placing passerines (< 20 g) and amphibian carcasses on stretches of roads we surveyed every 2 hours, we fine-tuned existing persistence estimates for these species. We found median disappearance times (time for half of the carcasses to disappear) of less than 30 minutes for birds and 1-18 hours for amphibians depending on the volume of traffic, which is much shorter than previous estimates. Rainfall did not impact carcass persistence. We show the implications of these results by (1) correcting the results of roadkill surveys performed on common toads in reproductive migration for the number of removed carcasses and (2) estimating the number of passerine-vehicle collisions in the Auvergne-Rhône-Alpes (AuRA) region of France using the citizen-science database Faune-AuRA. We estimate that a road survey conducted 3 hours after amphibian road crossing under-estimates the number of roadkill by half, and that about 6800 passerine collisions were missed in 2022 by contributors because of short carcass persistence on the road. Small-bodied carcasses are hard to detect for drivers, and for a collision-report rate of 1%, total collision numbers for passerines could be as high as 700 000 individuals from 21 reported passerine species in 2022 in AuRA.

## 1. Introduction

Human activities and their role in the on-going global biodiversity decline have been identified as the main cause of biodiversity loss over the past decades (Pollock et al., 2020). Of particular importance are linear transportation infrastructures such as roads (Forman et al., 2003) that threaten the viability of many wildlife populations (Moore et al., 2023), first and foremost of endangered species (Martin et al., 2018; Shepard et al., 2008; Taylor & Goldingay, 2009). The rapidly expanding road network concentrates all major threats of biodiversity at particular locations in space (Forman & Alexander, 1998; Rytwinski & Fahrig, 2015) jeopardizing the fitness of individuals. First, road surfaces hampers animal movement and gene flows through habitat fragmentation and destruction (Goosem, 2007; Lesbarrères & Fahrig, 2012; Prunier et al., 2014). Chemical pollutants accumulating on roads alter the composition of roadside vegetation communities, often favoring the expansion of invasive species (Lagerwerff & Specht, 2002; Nunes et al., 2020; Valladares et al., 2008). Traffic noise and light were shown to alter physiology and behavior, lower the reproductive success, and deter the presence of multiple species up to several hundreds of meters away from the road (Lengagne, 2008; Parris & Schneider, 2009; Reijnen & Foppen, 1994; Troïanowski et al., 2017). Finally, of all these ecological effects, roadkill is perhaps the most critical driver of biodiversity loss in relation to roads: wildlife-vehicle collisions most likely affect all terrestrial species, but on a scale we have yet to fully understand (Grilo et al., 2020, 2021; Moore et al., 2023).

One reason for this lack of knowledge is the quantification of roadkill numbers that are likely heavily underestimated due to unknown persistence and imperfect detection of dead animals along roads. Past studies suggested that performing road surveys on foot or bike at low speed improves accuracy by increasing the detection rates of roadkill carcasses (Barrientos et al., 2018; Ogletree & Mead, 2020). Once killed, an animal can either be scavenged, mechanically disappear with the flow of vehicles, or be displaced by people. This persistence time of carcasses on the road dictates the amount of time carcasses are at risk of detection (Ratton et al., 2014). The direct consequence of a persistence time of carcasses shorter than the frequency of surveys is a severe underestimation of roadkill numbers. Surprisingly, few authors have focused on the importance of carcass persistence and its structuring factors in roadkill surveys (but see Ratton et al., 2014; S. M. Santos et al., 2011; Teixeira, Coelho, Esperandio, & Kindel, 2013).

Most attempts at estimating the persistence time of carcasses focus on scavenging rates using chicks and other small-bodied animals as bait for carrion feeders, and find that theses carcasses seldom persist more than a few days after being placed on the road (Antworth et al., 2005; Ratton et al., 2014; R. A. L. Santos & Ascensão, 2019; Schwartz et al., 2018). Santos *et al*. (2011) recorded persistence times for a wide range of species and have highlighted the roles of road traffic, species and seasons on carcass removal rates. They found that carcasses disappeared quicker from smaller roads, possibly because low traffic volumes allow carrion-feeders easier access to roadkill (see also Ratton et al., 2014). Rain and high humidity also accelerated carcass disappearance. Humid conditions promote faster soft tissue decay (Brand et al., 2003), and Santos *et al*. (2011) suggested that in the case of roadkill, humidity promotes carcass dismemberment by vehicles. To our knowledge, there is no other published data on the relation between rain and roadkill persistence. The position of the carcass on the road also has been shown to influence persistence, with rodent carcasses on the lane disappearing faster than carcasses on the road shoulder (R. A. L. Santos & Ascensão, 2019).

The median persistence time (*i*.*e*., the time at which 50% of the carcasses have disappeared) of small-bodied fauna including reptiles, amphibians, small birds (<200 g), bats and other small mammals (<300 g) is approximately 1 day (R. A. L. Santos & Ascensão, 2019; S. M. Santos et al., 2011). Considering that in these studies, the surveys were done every 24 hours, median persistence time for these species might in reality be even less than a day. Indeed, Stewart (1971) placed 50 house sparrow carcasses on the surface of a highway and found that none remained after 2 hours. Roadkill surveys, even when performed daily, may under-estimate the number of collisions for the smallest species. Given the distribution of body size among mobile and terrestrial vertebrates (Smith & Lyons, 2013), road mortality could likely be underappreciated for 60% of the extent species. Consequently, efficient mitigation and conservation measures for these species require accurate estimations of persistence time and its spatio-temporal variability (Teixeira, Coelho, Esperandio, & Kindel, 2013).

Many small species of vertebrates are indeed of conservation concerns (Pereira et al., 2010). Roadkill data for small-bodied species such as rodents, small passerines or reptiles is scarce (but see Meek, 2009; Morelli et al., 2020; Ruiz-Capillas et al., 2015). Amphibian roadkill is better documented because massive roadkill occurs during the breeding season when amphibians leave their terrestrial habitat to gather in breeding ponds. It is the most endangered vertebrate group and has shown a significant decline in the last decades due to environmental disturbance (Wise, 2007). Mitigations measures were often used to reduce mortality during road crossings (Beebee, 2013; Helldin & Petrovan, 2019; Testud, 2020). Yet, for all small-bodied species including amphibians, we lack good quality data leading to robust estimations of persistence time, and consequently are left with unreliable estimates of roadkill occurrence.

To fill the gap in knowledge for small-bodied wildlife road mortality, we experimentally estimated the persistence time after a collision by placing carcasses on roads and monitoring their removal time and rate at a fine temporal resolution of 2 hours. We estimated persistence time on roads of different traffic volumes and under different weather conditions, predicting that persistence time would be shorter on smaller roads and in high humidity conditions (S. M. Santos et al., 2011). As carcass position on the lane could affect persistence (Antworth et al., 2005; R. A. L. Santos & Ascensão, 2019), we randomized the placement of the animal. Santos *et al*. (2011) found significant differences in persistence between species of similar body sizes but different taxa, so we used both small amphibians and birds (< 20g) in an effort to generalize our results to other small-sized species. Lastly, using roadkill surveys we performed during the reproductive migration of the common toad *Bufo bufo* and citizen-science data on small passerine road mortality, we estimated the proportion of missed carcasses, and therefore the number of amphibian roadkill during surveys as well as the number of small passerine roadkill in the Auvergne-Rhône-Alpes region France.

## 2. Material and Methods

### 2.1 Study location and sampling design

We measured persistence time between March and July 2022 for dry days, October and December 2022 for rainy days. We bought amphibian carcasses (*Pelophylax kl. esculentus)* from a licensed supplier (SONODIS) and procured small passerine carcasses (< 20g) from a wildlife rehabilitation center (Le Tichodrome, Isère, France). We stored all carcasses at -20°C and thawed them for 24h before the day of the experiment.

We selected 4 roads for passerine trials, and 7 roads for amphibians (speed limit: 80 km.h^-1^). We chose experimental locations to cover a wide range of road traffic volumes (130 to 9 300 veh.day^-1^). We retrieved the average traffic volume estimation over the last 3 years from local authorities for all roads but the smallest and less frequented ones (referred to as “*Petit Nice”* thereafter). We estimated missing traffic volume values by placing a camera-trap overlooking the road, and recording 1-minute videos every 5 minutes during 24 hours. We previously compared traffic estimations as provided by authorities and our method using camera traps, and found camera traps yielded very satisfactory results (camera-trap estimation: 11 220 veh.day^-1^; official mean daily traffic estimation: 11 135 veh.day^-1^).

On each road, we placed N=10 carcasses on the asphalt road surface. We chose the lane using a random number generator so that carcasses could be on road shoulders, in the center of the lane so they would fit in-between the tires of most vehicles, or directly in the path of the left or right vehicle tires (Fig. 2). We surveyed the carcasses on foot every two hours to determine whether they had disappeared. We considered a complete removal of a carcass when the observer could no longer detect it when walking on the road. Before running the experiment, we conducted training to ensure the different observers agreed on what a ‘removed’ carcass was. For passerines, we replicated the experiment on a rain-free day where the road surface was completely dry, and on a rainy day. For amphibians, we completed the experiments on the 8 locations during rain-free days but replicates on rainy days were limited to 3 locations only corresponding to traffic volumes of 130, 4160 and 8 000 veh.day^-1^.

### 2.2 Data analyses of persistence time

We obtained data in which the removal of the carcass from the road was only known to occur within an interval of two hours: for example, a carcass that was present on the first survey but had disappeared on the second had disappeared after a period of 2 to 4 hours on the road surface. The observations could be right-censored if the carcass was still present at the end of the experiment. As ignoring interval-censoring in survival data can lead to biases in survival estimates (Radke, 2003), we chose to model the persistence with interval-censored survival models, an extension of traditional survival models designed to analyze survival data in which the event of interest (here, disappearance from road) is only known to occur within a defined time interval (Anderson-Bergman, 2017). Interval censored models accommodate right-censored data points. We modeled the persistence of amphibians and small passerines in two separate models.

We used R package *icenReg* (Anderson-Bergman, 2017) for the R statistical software (R Core Team, 2021) which provides tools for implementing parametric interval-censored proportional hazards survival models. We selected the underlying parametric distributions of each model by visually comparing the available distributions to the corresponding Cox-PH model and selecting a distribution with no systematic deviation (Anderson-Bergman, 2017). Amphibian persistence was modeled using a gamma distribution, and passerine persistence using a log-normal distribution. We modeled the effects of road traffic volume (continuous variable), weather (2 modalities categorical variable: dry vs. rain) and position on the lane (2 modalities categorical variable: placed in the path of the tires or outside, see Fig. 2) on the persistence of the carcasses. We included observer ID (A/B) to accommodate for systematic differences in measurement between the two observers when analyzing amphibian data, but not for passerines because it lined up with the weather variable (all dry days were conducted by observer A, and all rainy days by observer B). We entered road traffic volume on persistence as a linear covariable, and as a log-transformed variable to account for potential non-linear relationships with carcass persistence. We represented the predicted median persistence by each model (linear vs. log-transformed) and the median persistence times of each road using the Turnbull estimator. We selected the log-transformed model for amphibian persistence and the linear model for passerine persistence based on visual fit (Fig. A1).

### 2.3 Estimation of roadkill numbers

Using our measurements of persistence times, we computed the number of roadkilled amphibians and small passerines during road surveys. For amphibians, we performed two roadkill surveys on foot on separate evenings in March 2022, during the common toad reproductive migration at a known road-crossing location on road D1504. The time elapsed between the collision window and the survey was 3 hours, and the D1504 road a mean daily traffic volume estimated at 3379 veh.day^-1^, which we adjusted to 1126 veh.day^-1^ considering the rule of thumb that only 25% of the traffic occurs between 7p.m. and 7a.m, the time of the survey, while the persistence experiments were conducted during the day (Van Langevelde & Jaarsma, 2005). Using the traffic volume and the time elapsed between collision window and road survey, we estimated from the survival model the proportion of carcasses having already disappeared from the road surface.

For passerines, roadkill count data was extracted for birds species < 20g from *Faune-Auvergne-Rhône-Alpes*, a citizen science database that compiles species presence as well as roadkill data submitted in real time by volunteers from the region Auvergne-Rhône-Alpes (thereafter: AuRA), France. We extracted the daily roadkill reports during the year 2022. Following Teixeira *et al*. (2012), the roadkill rate λ (collision.day^-1^) can be estimated as 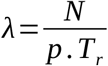 where *N* is the number of roadkill carcasses counted, *p* is the detection rate of the carcasses by the observer and the characteristic time *T*_*r*_ is the time needed to reduce the number of remaining carcasses on the road by ∽ 63%. We estimated *T*_*r*_ from the fitted survival model. We used it to estimate the daily number of collisions for 2022 in AuRA. We assumed detection rate *p* = 1 (all carcasses are detected) as we had no information about detection probabilities for small passerines. We explored the implications of imperfect carcass detection on passerine roadkill estimations in section 3 of the supplementary material.

To take into account the underlying distribution patterns of passerine species, we extracted the presence/absence data for each species in AuRA municipalities from Faune-AuRA, and for each municipality, computed the corresponding cumulative length of all roads accessible by car. We used this information to give estimates of the number of roadkill per kilometer of road in areas where the species was present.

## 3. Results

### 3.1 Carcasses persistence on the road

Half of the amphibians placed on the road disappeared within the first 5 hours on all roads except for the smallest (“*Petit Nice”* road, 130 veh.day^-1^). We projected that at t=24h, 95% of the amphibians should disappear from all roads, except for *Petit Nice* (52% at t=24h). In the following, we report model estimates for all covariables by hazard ratio (standard error); z-value and p-value. In accordance with our predictions, roads with higher traffic led to shorter amphibian carcass persistence: an increase of 50% in daily traffic increases the probability of removal from the road by 1.23 (hazard ratio = 1.67 (0.09); *z* = 5.62; *p* < 0.001; Fig. 1a). As predicted, we observed that probability of disappearance was 1.97 (0.23) times faster for amphibians placed in the path of car tires compared to road shoulder (z = 3.86; *p* < 0.001; Fig. 2a). However rain did not decrease or impact persistence (hazard ratio = 1.19 (0.3); *z* = 0.58; *p* = 0.56, Fig. 1a). There were no differences in amphibian persistence between observers (hazard ratio = 0.94 (0.29); *z* = -0.19; *p* = 0.85).

**Figure 1.**
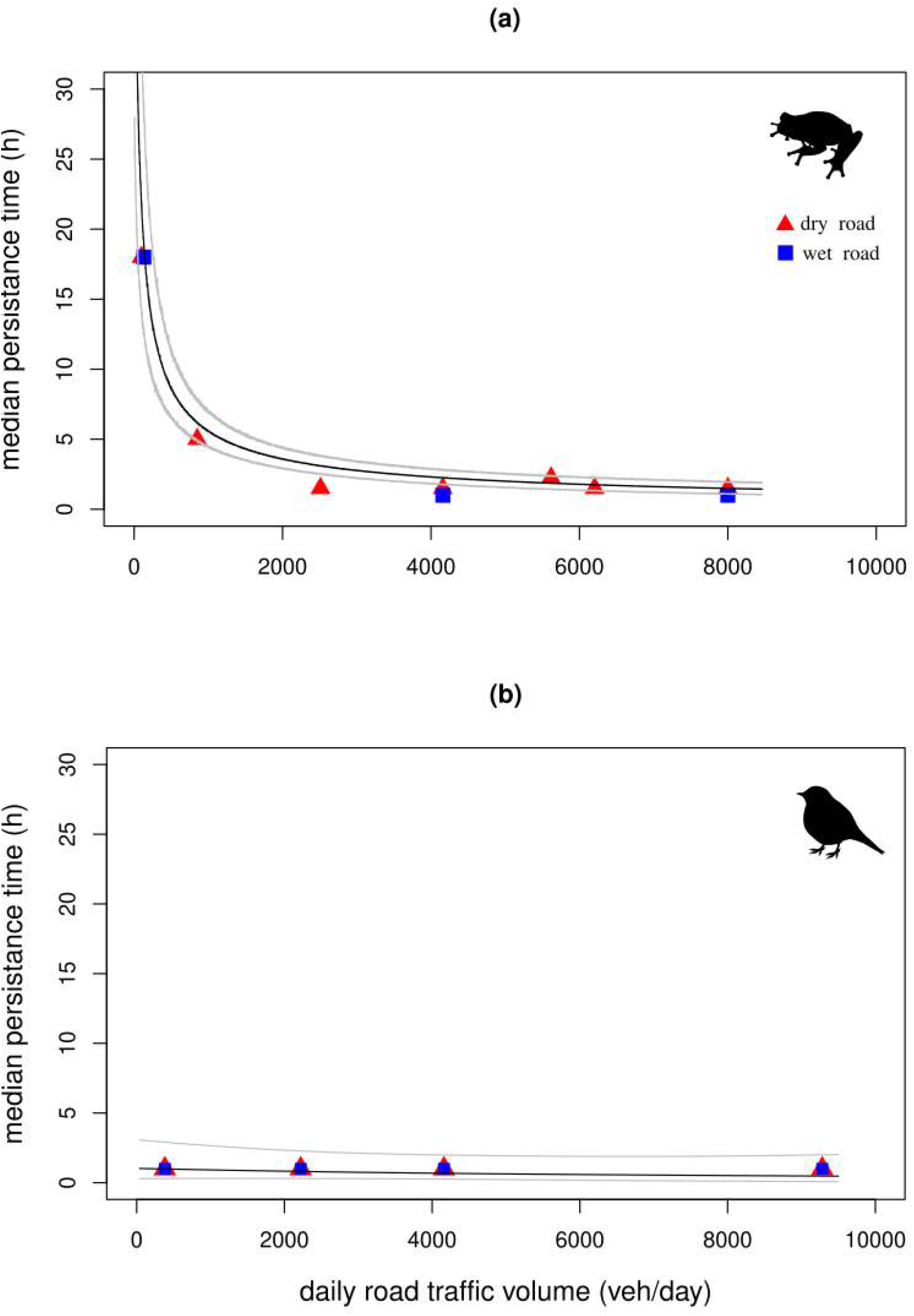
We performed roadkill persistence trials on 8 different roads for amphibians (a) and 4 roads for small passerines. At each location, 10 carcasses are placed on the road surface, and surveyed every two hours until complete disappearance. We represent the median persistence time (time for 50% of carcasses to disappear) of each road estimated during dry days without rainfall (▴) and rainy days (▪). Predictions from the fitted parametric survival models are represented as black lines (gray lines as the 95% confidence interval).

Within the first 2 hours, 80% of the passerines were no longer visible on any of the roads. We modeled road traffic volume as a linear covariable of persistence (Fig. A1). Our model predicted that virtually all passerine carcasses (97%) would disappear at time t = 24h. Contrary to all previous hypotheses, we found no differences in passerine persistence time depending on road traffic (hazard ratio = 1.0 (4.5e^-5^); *z*= 0.94 ; *p* = 0.347; Fig. 1b), carcass position on the lane (hazard ratio = 0.99 (0.27); *z* = -0.003; *p* = 0.996; Fig. 2b) or weather (hazard ratio = 1.23 (0.26); *z* = 0.81; *p* = 0.420; Fig. 1b).

**Figure 2.**
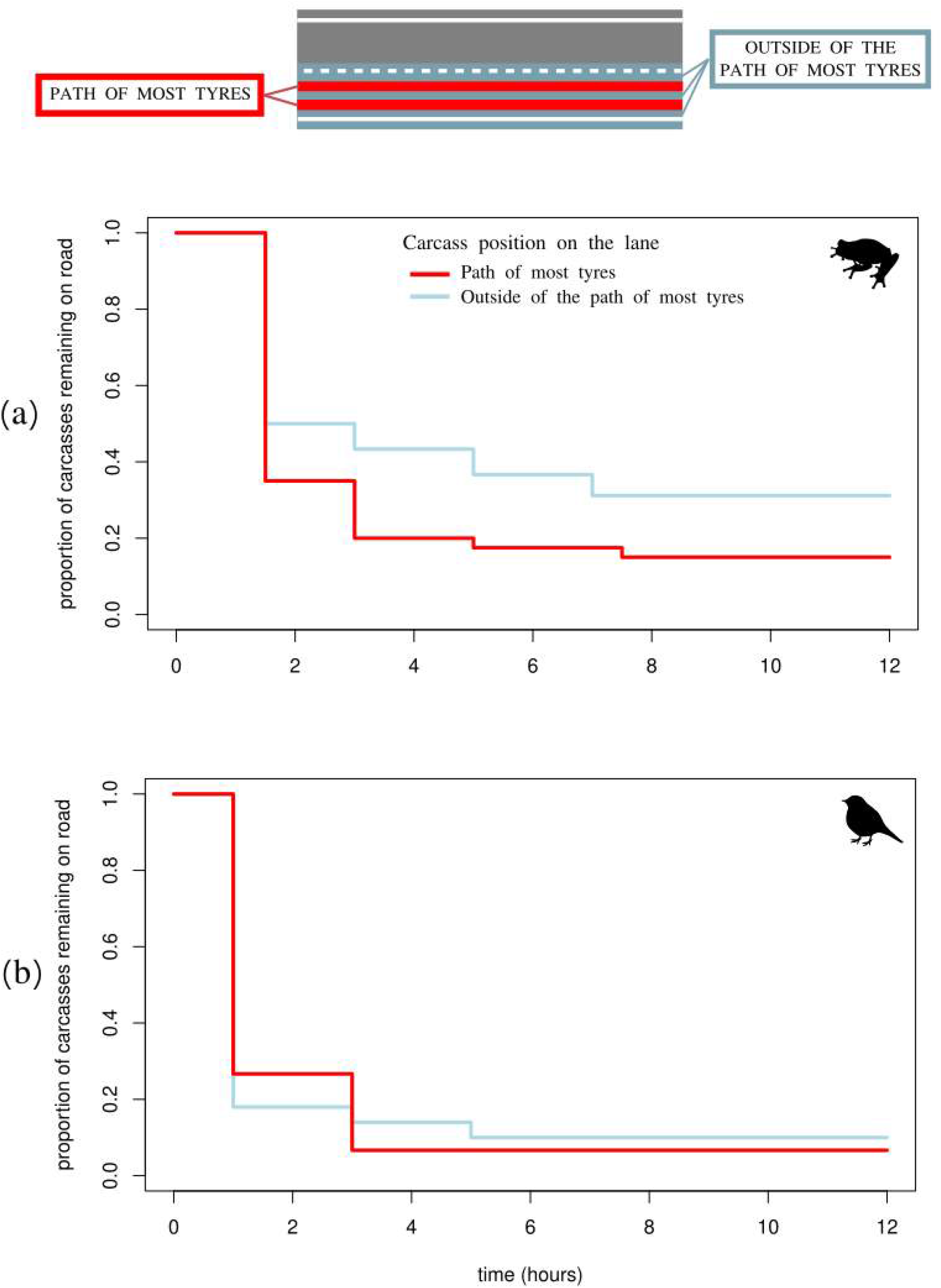
We randomly positioned carcasses of 90 frogs (a) and 80 carcasses from multiple species of small passerines (b) on the surface of different roads. We represent here the Kaplan-Meier survival curves of the first 12 hours of carcass persistence time across all roads. Carcasses were randomly positioned to be either in the path of the tires of most 4-wheeled vehicles (red), or outside of the path of the tires (blue). Parametric survival models showed that frog carcasses disappeared faster when placed in the path of vehicle tires while passerines had similar persistence on every position on the lane.

### 3.2 Practical implications for roadkill estimations

On March 15^th^ and March 20^th^ 2022, we counted respectively 12 and 30 common toad carcasses on road D1504. Using the fitted survival model of amphibian persistence described above, we predicted the proportion of missing carcasses on the road according to traffic volume. After 3 hours, we estimated that 49.6% (95% confidence interval: 39.4, 60.5) of the carcasses had already been removed from the road at the time of survey meaning that, assuming perfect detection of carcasses on the road, they were 17.9 (16.7, 19.1) toad casualties on March 15^th^ and 44.8 (41.8, 48.1) on March 20^th^.

For passerines, 201 small passerines from 21 species were reported by contributors in 2022 (see Table 1 for a complete list). We estimate the characteristic time T_r_ at 0.0283 days (95% confidence interval: 0.009, 0.088). Accounting for carcass removal rates amounts to 7093 (2270, 22120) passerines roadkilled in 2022 in the AuRA region (Table 1). The most commonly killed species on roads is the house sparrow *Passer domesticus* with 0.87 individuals killed per year every 100km and the least common is the long-tailed tit *Aegithalos caudatus* with 0.019 individuals. These estimates assume that carcasses are always detected and reported by Faune-AuRA contributors. Straightforward estimates of roadkill numbers accounting for imperfect detection and report can be derived from these results: assuming that 1% of passerine carcasses present on the road are seen and reported, 0.87/0.01 = 87 house sparrows and 0.019/0.01 = 19 long-tailed tits are killed annually per 100km of road, and 709 300 (227 000, 2 212 000) passerines could have been killed on roads in the AuRA region in 2022 (Fig. A2).

**Table 1:** We used roadkill reports for small passerine (< 20g) in the Auvergne-Rhône-Alpes (AuRA) region of France from the citizen-science database Faune-AuRA in 2022. We corrected the reports for carcass removal rate from road (Teixeira *et* al., 2012) and then, using the presence data from Faune-AuRA to infer species distribution patterns, we give estimations of roadkill per 100 km of road in areas where the species is present. These estimations of roadkill rates account for the speed of carcass disappearance from the road but not for the number of carcasses missed or not reported by the observers.

|  | Reported in<br>Faune-AuRA<br>(2022) | Number of collisions after<br>correcting for persistence<br>(95% confidence interval) | Number of roadkill<br>per 100km of road<br>(95% confidence interval) |
| --- | --- | --- | --- |
| Long-tailed tit<br>*Aegithalos caudatus* | 1 | 35.3 (11.3, 110.1) | 0.019 (0.006, 0.06) |
| European goldfinch<br>*Carduelis carduelis* | 5 | 176.5 (56.5, 550.3) | 0.081 (0.03, 0.25) |
| Eurasian blue tit<br>*Cyanistes caeruleus* | 2 | 70.6 (22.6, 220.1) | 0.030 (0.01, 0.09) |
| Cirl bunting<br>*Emberiza cirlus* | 1 | 35.3 (11.3, 110.1) | 0.025 (0.008, 0.08) |
| Yellowhammer<br>*Emberiza citrinella* | 2 | 70.6 (22.6, 220.1) | 0.069 (0.022, 0.21) |
| European robin<br>*Erithacus rubecula* | 58 | 2046.9 (655.2, 6382.9) | 0.84 (0.28, 2.63) |
| Common chaffinch<br>*Fringilla coelebs* | 16 | 564.6 (180.7, 1760.8) | 0.23 (0.07, 0.7) |
| White wagtail<br>*Motacilla alba* | 6 | 211.8 (67.8, 660.3) | 0.12 (0.03, 0.33) |
| Great tit<br>*Parus major* | 10 | 352.9 (112.9, 1100.5) | 0.14 (0.05, 0.44) |
| House sparrow<br>*Passer domesticus* | 56 | 1976.3 (632.6, 6162.8) | 0.87 (0.28, 2.7) |
| Black redstart<br>*Phoenicurus ochruros* | 11 | 388.2 (124.3, 1210.6) | 0.18 (0.06, 0.56) |
| Common redstart<br>*Phoenicurus phoenicurus* | 3 | 105.9 (33.9, 330.2) | 0.081 (0.03, 0.25) |
| Willow warbler<br>*Phylloscopus trochilus* | 1 | 35.3 (11.3, 110.1) | 0.05 (0.02, 0.15) |
| Whinchat<br>*Saxicola rubetra* | 1 | 35.3 (11.3, 110.1) | 0.052 (0.02, 0.16) |
| European stonechat<br>*Saxicola rubicola* | 6 | 211.8 (67.8, 660.3) | 0.13 (0.04, 0.42) |
| European serin<br>*Serinus serinus* | 1 | 35.3 (11.3, 110.1) | 0.022 (0.007, 0.07) |
| Eurasian blackcap<br>*Sylvia atricapilla* | 12 | 423.5 (135.6, 1320.6) | 0.18 (0.06, 0.56) |
| Garden warbler<br>*Sylvia borin* | 1 | 35.3 (11.3, 110.1) | 0.057 (0.02, 0.18) |
| Common whitethroat<br>*Sylvia communis* | 1 | 35.3 (11.3, 110.1) | 0.034 (0.01, 0.11) |
| Sardinian warbler<br>*Sylvia melanocephala* | 1 | 35.3 (11.3, 110.1) | 0.14 (0.04, 0.44) |
| Eurasian wren<br>*Troglodytes troglodytes* | 3 | 105.9 (33.9, 330.2) | 0.05 (0.02, 0.16) |
| **TOTAL** | **201** | **7093 (2270.6, 22120.2)** | **3.39 (1.08, 10.57)** |

## 4. Discussion

### 4.1 Persistence on road and implications for road monitoring

Most small carcasses of amphibians and birds in this experiment were removed from the road surface in less than a day. We estimated the median persistence time (*i.e*., the time needed for half of the carcasses to be removed from the road) of small passerines (< 20g) at *t =* 22 minutes using a parametric survival model, and this persistence was not affected by road traffic volume, position of the carcass on the road or rain. Common toad carcass persistence was similarly unaffected by rain but was largely dependent on the traffic volume of the road within the explored range (130 - 9000 veh.day^-1^): the more vehicles, the less time the carcasses remained visible. On roads with 1000 vehicles per day or more, amphibian median persistence time is 5 hours or less. Amphibians placed directly in the path of the tires disappeared nearly twice as fast than the amphibians positioned elsewhere on the road surface.

Santos *et al*. (2011) found that 50% of small carcasses disappeared in one day for small-sized species (amphibians, reptiles and small birds). Our results agree with these previous observations, and the fine-tuning of previous estimations shows that the median persistence time is actually less than 24h for both amphibians and small birds. For passerines, 80% of carcasses had disappeared within 2 hours across all locations, and 94% after 12h. For amphibians, after 12h we found that over half of the carcasses had disappeared across all locations: 60% had disappeared from *Petit Nice*, a road with little vehicular traffic, and none remained on the major road *Caluire Saône*. Daily roadkill surveys have been previously recommended for accurate roadkill data for small-bodied species (Henry et al., 2021; S. M. Santos et al., 2015). Our results suggest instead that even daily road monitoring will largely underestimate the actual number of collisions of amphibians and small passerines with road vehicles. In fact, the short persistence of these species on the road indicates that roadkill surveys will always underestimate casualties unless we monitor roads continuously. Because this is an impossible task, authors should focus instead on implementing persistence corrections (see Teixeira, Coelho, Esperandio, & Kindel, 2013) to their survey methodology. Body size is often cited as the biggest predictor of carcass persistence time on the road (Barrientos et al., 2018; Guinard et al., 2012; Henry et al., 2021). Nevertheless, we found substantial differences in the persistence time between amphibians and small passerines of similar sizes. Hence, the generalization of persistence times between small-bodied species is not possible, and our results are not suitable to assess the persistence of lizards, snakes, bats and other smalls mammals. We suggest instead that other experiments of persistence with fine-tuning should be conducted on these species to assess the proportion of missed carcasses by daily road surveys.

Scavenger activity is often cited as the main cause of carcasses disappearance in roadkill persistence studies (Antworth et al., 2005; Erritzoe et al., 2003; Slater, 2002). For instance, Schwartz et al (2018) determined that fresh roadkill attracts corvids and foxes that could remove up to 62% of the carcasses within 2 hours. Smaller roads might be easier to access for carrion-feeders than highly-frequented ones, and roadkill persistence studies report that roadkill generally disappeared faster on smaller roads compared to more frequented ones (S. M. Santos et al., 2011; Slater, 2002).We found instead that amphibian persistence decreased with the volume of traffic, suggesting that this is not true for all taxa, or that scavenging is not the main mechanism of carcass removal in this case. It has been suggested that common toads are not attractive to carrion-feeders due to their though skin (Hels & Buchwald, 2001). Instead, mechanical fragmentation of tissues by cars appears to play a major role in amphibian carcass disappearance: the accumulation of repeated crushing of biological tissues by tires flattened most of the carcasses we placed on the road surface that became less and less visible at each observation. This mechanism is consistent with findings that traffic volume significantly decreases persistence time in amphibians (Fig. 1a) and that carcasses directly in the path of the tires for most vehicles were removed 2 times faster compared to carcasses placed outside of it (Fig. 2a).

Passerine carcasses disappeared at higher rates than amphibians (80% within the first 2 hours) and at the same speed across all tested locations and all positions on the lane. Ratton *et al*. (2014) and Antworth *et al*. (2005) found longer carcass persistence when using *Gallus domesticus* chick carcasses (resp. 66% removed within 12h and 40% within 2h). However, the chick carcasses weighted slightly more than the wild passerine species used here: reportedly around 30 grams in both studies, against 20g or less in this experiment. Domestic species are also known to have higher removal rates than wild species in open fields (Prosser et al., 2008). To our knowledge, the only other roadkill persistence experiment at a fine temporal resolution using passerines was conducted by Stewart (1971) with house sparrows *Passer domesticus*. Interestingly, he reported persistence estimates similar to our results (100% removal within the first two hours) on a highway but found very contrasted results on a smaller country road: all carcasses were still visible after 12 hours. On the contrary, and similarly to our findings, chick persistence did not vary between a frequented highway and a low traffic dirt road in Ratton *et al*. (2014). Small birds persistence on roads appear to be highly variable between species and locations.

Contrary to common toads, passerine persistence on roads showed no correlation to road traffic volume. It could be that the temporal resolution of 2-hours intervals is not fine enough to capture the variability in passerine removal rates from roads. During the experiments, and contrary to amphibians, bird carcasses often disappeared without a trace in-between surveys, which could instead suggest that the high removal rate is the result of scavenging activity (Antworth et al., 2005). We conducted a complementary experiment with continuous observation of the carcasses during 2 hours (Supp. Mat. Section 3). We placed 12 passerine carcasses randomly on a highly-frequented portion of road (8000 veh.day^-1^) in an urban setting with no unpaved shoulder where the speed limit was 50 km.h^-1^. We found a higher persistence than previous estimates : 5 out of 12 (41%) were invisible by the end of the 2 hours, against an average across all roads of 80% in the main experiments. Carcasses disappeared at variable rates, with some becoming undetectable after the passage of 10 vehicles and other being still visible after 800 vehicles (median = 178 vehicles, Fig. A3). All carcasses where crushed by vehicle’s tires, including carcasses originally placed out of the path of the tires (center of the road, center of the lane in-between tires, roadside) that were either crushed by vehicles swerving on the lane to avoid obstacles such as parked cars or displaced by the turbulence following passing vehicles. Contrary to the common toad, we observed that passerine body shape and feathers could promote carcass mobility on the road surface. Passerines carcasses that are displaced by passing vehicles can also potentially end up in the vegetation of the road shoulder, masking them from view during surveys (AB personal observations).

Season and weather influence the speed of carcass disappearance in open fields where removal is usually the result of decomposition or scavenging, and the highest rates of removal are found during the summer months where temperatures are high (Costantini et al., 2017; Prosser et al., 2008; Sharanowski et al., 2008). Carcass persistence on roads was instead found to be lower during spring in France (Guinard et al., 2012), and high temperatures and humid conditions accelerated the disappearance of carcasses from roads in Portugal (S. M. Santos et al., 2011). We tested here the effect of rainfall on persistence time, and expected carcasses to disappear faster under the rain. We found instead similar persistence between dry and rainy days for both amphibians and passerines (Fig. 1). Rain and humidity might not make a difference when a small-bodied fresh carcass is confronted to a tire, as opposed to bigger animals. Instead of weather, persistence on the road for small species could vary between seasons following scavenger activity (Guinard et al., 2012; Schwartz et al., 2018).

### 4.2 Impact of vehicular collisions in biodiversity loss

Collisions of wildlife with vehicles have long-been suspected of being a significant threat to the persistence of many amphibian populations (Glista et al., 2008; Puky, 2005) but estimating the biodiversity loss in terms of abundance remains a major challenge. In the case of common toads, because amphibian roadkill during the reproductive migration happens during a short time window, we were able to estimate the time since collision. When the speed of carcass removal from the road is also known, estimates of the proportion of carcasses that have disappeared and were subsequently missed during the survey are straightforward, assuming that all toads present on the road during the survey were detected. In the roadkill survey we conducted, ignoring carcass persistence leads to underestimations of the number of toad roadkill of about -50%, for a survey conducted 3 hours after amphibian crossing at this particular location. A similar survey conducted the next morning (t=12h after road crossing window) would have under-estimated the casualties by about 70%. Aquatic breeding amphibians are particularly suited to this example of roadkill counts correction because roadkill is synchronized at sunset, allowing for estimations of the time elapsed between the road crossings and the survey, and because road crossings are localized in space, allowing for on-foot surveys where the detection rate of carcasses is higher than surveys done by car.

Correcting roadkill estimates for carcass persistence is much less straightforward when animal-vehicle collisions happen continuously. Teixeira *et al*. (2013) developed a theoretical framework of carcass persistence correction for roadkill monitoring surveys which we applied to citizen-science data of passerine roadkill. We estimated a total of 7000 collisions between small passerines and vehicles in the AuRA region. Not all small passerine species present in AuRA were present in the roadkill reports for 2022, meaning that this number is likely under-estimated. Roadkill rates ranged from 0.9 (house sparrow *Passer domesticus* and european robin *Erithacus rubecula)* to 0.01 (long-tailed tit *Aegithalos caudatus)* collision per 100km of road and per year. Roadkill rates are likely linked to the local abundance of species (Seiler & Helldin, 2006). Despite having no current estimate of these species’ abundance in the AuRA region, *P. domesticus* and *E. rubecula* are the most reported small passerines in Faune-AuRA, and they are reported 5 times more than *A. caudatus* (resp. 562 577, 681 000 and 164 600 direct observations of living specimens between 2000 and 2023). This shows that patterns of species abundance in the AuRA region could be a driving factor in the inter-specific differences in roadkill rates, along with species traits and behaviors (Grilo et al., 2020; Teixeira et al., 2013).

However, while we estimate the persistence time on the road for these species, we have little information about the conversion rate of roadkill carcasses into reported roadkill. Indeed, following a collision with a vehicle a passerine might never be detected by contributors (detection rate) or be detected but never reported (reporting rate), because they forgot or could not identify the species with certainty. Additionally, an unknown proportion of passerines could be projected into the roadside vegetation at the moment of impact. Very little work has been done on roadkill detection, although we know that detection rates of roadkill from a car (which encompasses the majority of reports into Faune-AuRA) is significantly lower than when surveying carcasses on foot (Teixeira et al., 2013). Reporting rates in citizen-science roadkill projects are even less documented. We estimated the cumulative number passerine roadkill for the 21 reported species using a range of plausible values for the detection and reporting probabilities in figure A2. Using an estimate of 1% of report for passerine collisions with vehicles (*i.e*., 1% of the passerines killed by a vehicle are detected, identified and reported by a contributor) and after correcting for passerine persistence time on the road, we estimate about 700 000 small passerines killed by vehicles in 2022 in the AuRA region, a surface of 70 000 km². In other words, the roadkill citizen-science reports only give us a glimpse into a much larger issue, as we multiplied the number of reports by a factor of 3500. Currently, no estimates of the abundance of passerine species in the region are available to put these estimates into perspective. The French Breeding Bird Survey (a standardized protocol for bird populations tracking, Julliard & Jiguet, 2002) reports alarming declines of up to -50% in wild small passerine french populations over the last two decades (european greenfinch *Chloris chloris*, -50%; european serin *European serin*, -41.7%; yellowhammer *Emberiza citrinella, -*53.6%). Although wild passerine populations face multiple anthropogenic threats from farmland practices (Broyer et al., 2016; Moreau et al., 2021; Rigal et al., 2023), road mortality could be an important and under-estimated driver of biodiversity loss that needs investigating.

## Conclusion

By conducting daily or bi-daily surveys, an observer would miss an important part of the road mortality for small-bodied birds and amphibians. Adapting the periodicity of roadkill surveys to minimize the number of missed carcasses is not a viable option (unless we monitor road continuously). Instead, we should focus on quantifying the persistence of carcasses and correct the data gathered during surveys. Our estimations of the number of killed animals by collisions with vehicles illustrate the huge gap between raw counts of carcasses on roads and how many animals were actually lost. Common toads casualties 3 hours after they crossed the road were already underestimated by half, which has severe consequences in the quantification of the impact of road mortality in amphibian biodiversity loss. Assuming a collision-report rate of 1% in the citizen-science project *Faune-Auvergne-Rhône-Alpes*, we estimate 700 000 individuals killed each year in a surface of 70 000 km², which could play a part in the national decline in small passerine population observed over the last two decades.

## Acknowledgments

We thank the wildlife rehabilitation center Le Tichodrome for their help in providing small passerine carcasses and Benoit Vallas for his contribution to the data record.

## Supplementary material

**Figure A1:**
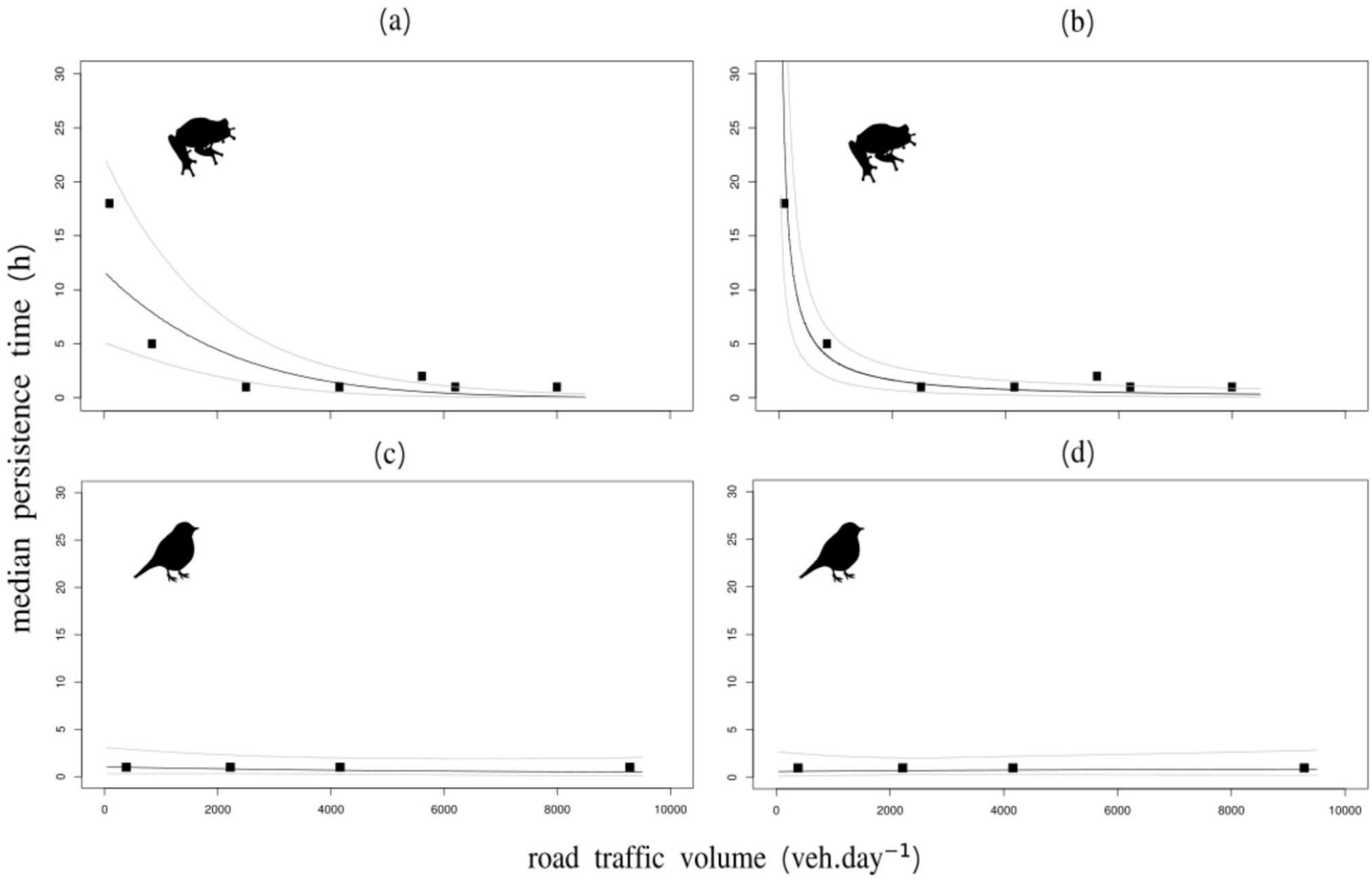
We represent the median persistence time (▪; time for half of the carcasses to disappear) of amphibian (a, b) and small passerine (c, d) carcasses on roads of different volumes of traffic. We fitted parametric survival models for both taxa, including road traffic volume as one of the covariables. The predictions of the models are represented as black lines (gray lines: 95% confidence interval). Road traffic volume was either modeled as a linear (a, c) or log-linear (b, d) covariable, and we selected the best option based on the visual fit between the median persistence on each road and the model predictions. Amphibian persistence was modeled using a log-linear effect of traffic (b) and passerine persistence was modeled using a linear effect (c).

**Figure A2:**
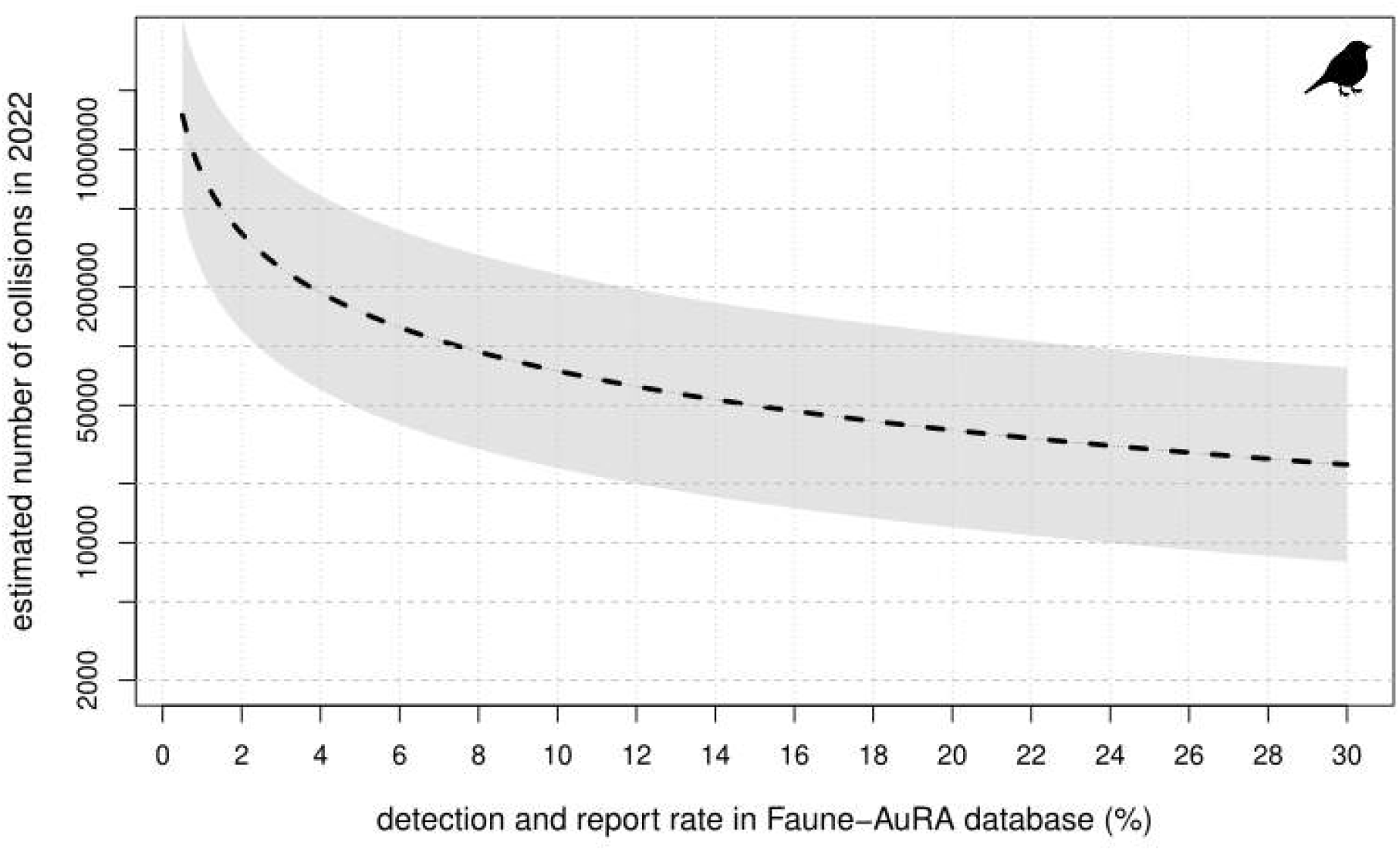
Estimated number of collisions (gray area: 95% confidence interval) between 21 small passerine species (< 20g) and road vehicles in 2022 in the region Auvergne-Rhône-Alpes (AuRA), France, based on reports from citizen-science database Faune-AuRA. Roadkill reports were first adjusted to account for carcass persistence time on the road (tested experimentally with fresh carcasses) following Teixeira *et al*. (2013). Then, following the assumption that only a small proportion of small-bodied carcasses present on the road are both seen and reported by users in citizen-science roadkill reporting projects, we show here the estimated number of actual collisions if between 0.5 and 30% of passerines carcasses present on the road surface end up in the database. For example, if 1% of the visible small passerine roadkill is reported, we estimate 709 100 actual collisions in AuRA in 2022. Complete list of small passerines species reported in Faune-AuRA in 2022: *Aegithalos caudatus, Carduelis carduelis, Carduelis chloris, Cyanistes caeruleus, Emberiza cirlus, Emberiza citrinella, Erithacus rubecula, Fringilla coelebs, Fringilla montifringilla, Motacilla alba, Motacilla flava flava, Parus major, Passer domesticus, Phoenicurus ochruros, Phoenicurus phoenicurus, Phylloscopus trochilus, Prunella modularis, Saxicola rubetra, Saxicola rubicola, Serinus serinus, Sylvia atricapilla, Sylvia borin, Sylvia communis, Sylvia melanocephala, Troglodytes troglodytes*.

**Figure A3:**
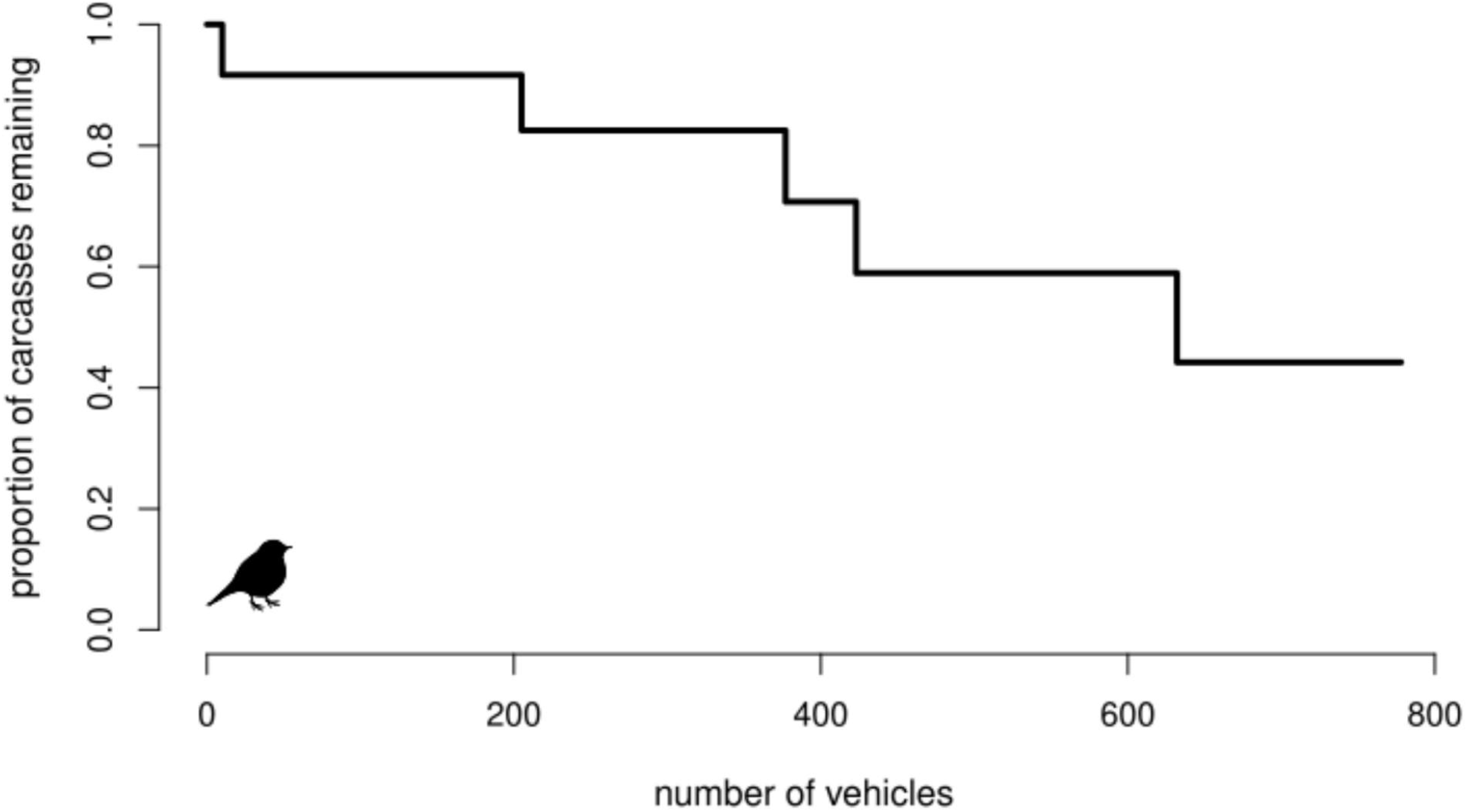
We conducted an additional experiment where we placed 12 small passerine carcasses on a dry road surface (8000 veh.day^-1^) and continuously counted the number of passing vehicles until the carcasses disappeared (*i.e*. carcass can’t be detected on foot during a road survey or species is no longer identifiable). The experiment was conducted on an urban road with a 50 km.h^-1^ speed limit, during a day with no rainfall. Passerines were placed randomly on the road surface using a random number generator. Half of the carcasses had disappeared after 632 vehicles, and 58% were still visible when the experiment was ended after 2 hours. All carcass disappearances were the result of repeated crushing by vehicle wheels. All birds (100%) were crushed at least once during the experiment, including carcasses placed in parts of the road lane that vehicles were unlikely to reach, either because the turbulence of passing vehicles had displaced the carcasses, or because some vehicles swerved on the lane to avoid obstacles. The rates of carcass removal during the first 2 hours are lower here than in the experiments conducted in the main body of this article, which hints at a possible involvement of other mechanisms of carcass removal beyond mechanical crushing by tires.

